# Gene expression response to sea lice in Atlantic salmon skin: an RNA-Seq comparison between resistant and susceptible animals

**DOI:** 10.1101/225094

**Authors:** Diego Robledo, Alejandro P. Gutiérrez, Agustín Barría, José M. Yáñez, Ross D. Houston

## Abstract

**Background:** Sea lice are parasitic copepods that cause large economic losses to salmon aquaculture worldwide. Frequent chemotherapeutic treatments are typically required to control this parasite, and alternative measures such as breeding for improved host resistance are desirable. Insight into the host-parasite interaction and mechanisms of host resistance can lead to improvements in selective breeding, and potentially novel treatment targets. In this study, RNA sequencing was used to study the skin transcriptome of Atlantic salmon parasitized with sea lice (*C. rogercresseyi*). The overall aims were to compare the transcriptomic profile of skin at louse attachment sites and ‘healthy’ skin, and to assess differences between animals with varying levels of resistance to the parasite.

**Results:** Atlantic salmon were challenged with *C. rogercresseyi*, growth and lice count measurements were taken for each fish. 21 animals were selected and RNA-Seq was performed on skin from a louse attachment site, and skin distal to attachment sites for each animal. These animals were classified into family-balanced groups according to the traits of resistance (high vs low lice count), and growth during infestation (an indication of tolerance). Overall comparison of skin from louse attachment sites versus healthy skin showed that 4,355 genes were differentially expressed, indicating local up-regulation of several immune pathways and activation of tissue repair mechanisms. Comparison between resistant and susceptible animals highlighted expression differences in several immune response and pattern recognition genes, and also myogenic and iron availability factors. Genomic regions showing signs of differentiation between resistant and susceptible fish were identified using an Fst analysis.

**Conclusions:** Comparison of the skin transcriptome between louse attachment sites and healthy skin has yielded a detailed profile of genes and pathways with putative roles in the local host immune response to *C. rogercresseyi*. The difference in skin gene expression profile between resistant and susceptible animals led to the identification of several immune and myogenic pathways potentially involved in host resistance. Components of these pathways may be targets for studies aimed at improved or novel treatment strategies, or to prioritise candidate functional polymorphisms to enhance genomic selection for host resistance in commercial salmon farming.

## BACKGROUND

Aquaculture is currently the fastest growing food industry [1] and is essential to meet increasing global demands for fish. However, the sustainability and prolonged success of any farming industry depends on effective disease prevention and control, and this tends to be particularly challenging for aquaculture. The aquatic environment and high stock density can expedite pathogen spread, which has historically resulted in periodic mass mortality events and ongoing challenges in disease prevention and control. While biosecurity measures, vaccination, nutrition and medicines all play vital roles for several diseases, selective breeding to produce more resistant and tolerant aquaculture stocks is rapidly becoming a key component of the battle to prevent these outbreaks [2, 3].

Sea lice, ectoparasites of the family Caligidae, are one of the major disease problems facing the aquaculture industry, and specifically for salmon farming. Atlantic salmon is the most important species in aquaculture with a production value of 14.7 billion US dollars in 2014 [1], therefore control of sea lice is a primary goal for the industry. Sea lice-related economic losses to worldwide salmonid aquaculture were estimated at ~430 million USD per annum [4]. Two lice species present the primary concerns for salmon farming: *Lepeophtheirus salmonis* in the Northern Hemisphere and *Caligus rogercresseyi* in the Southern [5]. These copepods parasitize salmon during the marine phase of the lifecycle by attaching to their skin or fins, and feeding on the blood and tissue. This leads to open wounds which can facilitate the entry of other pathogens. The impaired growth and secondary infections cause significant negative animal welfare and economic impact [6]. Interestingly, the outcome of sea lice infestation varies for different salmonid species, with coho salmon (*Oncorhynchus kisutch*) showing rapid inflammatory response and epithelial hyperplasia, leading to parasite encapsulation and more than 90% reduction in lice loads [7]. In comparison, Atlantic salmon (*Salmo salar*) is highly susceptible to sea lice infestation and seemingly cannot mount an effective immune response [7].

Despite extensive use of both chemical and non-chemical treatments to control sea lice, their negative impact on salmon aquaculture has increased in the past years [8]. Various sea lice populations have been reported to be resistant to the most common chemicals available for therapeutic control [9]. Therefore, alternative methods to control sea lice are currently being studied, including the use of probiotics to reduce salmon attractiveness for sea lice [10] or cohabitation with lice-eating species [11, 12]. A promising and potentially complementary approach to existing control measures is to exploit natural genetic variation in farmed salmon populations to breed stocks with enhanced resistance to the parasite. Current evidence indicates that host resistance to sea lice in Atlantic salmon has a highly polygenic genetic component [13, 14, 15, 16, 17]. The presence of significant genetic variation for resistance against *Caligus royercresseyi*, with heritability values ranging between 0.1 up to 0.34, demonstrates the feasibility of improving this trait by selective breeding in Atlantic salmon [18, 19]. While selective breeding for resistance can be effective without any prior knowledge of the underlying genes, understanding the functional basis underpinning genetic resistance is a major research goal. The benefits of this include the prioritisation and testing of putative functional (causative) variants which may be directly affecting host resistance, for which the new Functional Annotation of All Salmonid Genomes (FAASG) project will be integral [20]. Such variants can enhance selective breeding by improving accuracy of prediction of the genetic merit, particularly across distant relatives or populations. Finally, these causative variants could be introduced into populations or species where it has never been present through the use of genome editing, for example using CRISPR-Cas9 technology which has been successfully applied in salmon [21].

In addition to the value to selective breeding programs, knowledge of the interaction between salmon and sea lice can help devise more effective prevention and treatment strategies. RNA sequencing can provide a relatively holistic view of the host response to parasite infection, which in turn can highlight specific genes, pathways and networks involved in the host-parasite interaction. RNA-Sequencing can also be used to identify genetic markers in transcribed regions, and to assess the putative impact of those markers on transcript and protein function. The effect of these markers on gene expression (and ultimately host resistance) can be assessed, leading to a shortlist of candidate functional variants. The overall aims of the current study were to compare the transcriptome profile of salmon skin at louse attachment sites and ‘healthy’ skin, and to evaluate differences in these profiles between animals with varying levels of resistance to the parasite. To achieve this, challenged animals were classified into family-balanced groups according to the traits of resistance (based on high vs low lice count), and growth during infestation (an indication of tolerance) and RNA-Seq was performed on individual samples. While individual growth rate during challenge is not strictly ‘tolerance’ (because it is not scaled according to parasite burden [22]), this is the term used herein to refer to this trait. By comparing these groups, genes, pathways and genetic variation related to local immune response and host resistance were identified and discussed.

## RESULTS AND DISCUSSION

### Disease challenge

A total of 2,632 fish belonging to 105 families from a commercial breeding program were challenged with *Caligus rogercresseyi*, and killed for sampling eight days post challenge. Average lice burden per fish was 38 ± 16. Fish were selected for RNA sequencing based on the traits of resistance, measured as number and concentration of lice per fish, and tolerance, defined in this study as weight and length gain since the start of the challenge. The selected fish allowed for 8 vs 8 comparisons between family-matched fish showing differential resistance (26.2 ± 5.5 vs 54.9 ± 13.5 sea lice per fish) and differential tolerance (7.0 ± 4.3 vs 28.8 ± 12.3 weight gain percentage). A total of 42 samples (21 fish, skin from sites of louse attachment and healthy skin) were sequenced, resulting in an average of ~27.9 ± 2.7 million reads per sample. After trimming, these were aligned against the salmon reference genome (ICSASG_v2; Genbank Accession GCF_000233375.1 [23]) and levels of gene expression were estimated according to the official salmon genome annotation (NCBI *Salmo salar* Annotation Release 100). 25% of the aligned reads were discarded due to multimapping, and the position of an additional 11% did not overlap with any known gene. Following these filters, an average of 19 M reads per sample were assigned to genes and used for downstream analyses of gene expression. All raw sequence data is available in NCBI’s Sequence Read Archive (SRA) under BioProject accession number SRP100978, and may be a useful contribution to the functional annotation of all salmonid genomes initiative (FAASG [20]).

### Louse attachment sites versus healthy skin

Hierarchical clustering of samples according to their normalized gene expression did not initially reveal an obvious clustering according to any of the measured phenotypes, nor between samples from louse attachment sites and those from healthy skin (Additional file 1). Despite this, after a single outlier sample was removed, a total of 1,711 differentially expressed (DE) genes were identified between healthy and injured skin (Figure 1A). These DE genes showed relatively small log_2_ fold change (FC) differences. The expression levels for the most significant DE genes from this list (p < 0.001, 156 genes) are shown in a heatmap (Figure 1B), which highlights a gene expression signature of lice attachment sites and healthy skin. However, a small number of lice attachment site samples did not cluster as expected and were removed. The differential expression was repeated using 13 vs 13 samples (Figure 1C). Despite the lower statistical power due to the reduced sample size, the number of significant DE genes was substantially larger (n = 4,355, Additional file 2), with a higher number of up-regulated than down-regulated genes when comparing attachment sites to healthy skin (n = 3,114 vs n = 1,241).

**Figure 1.**
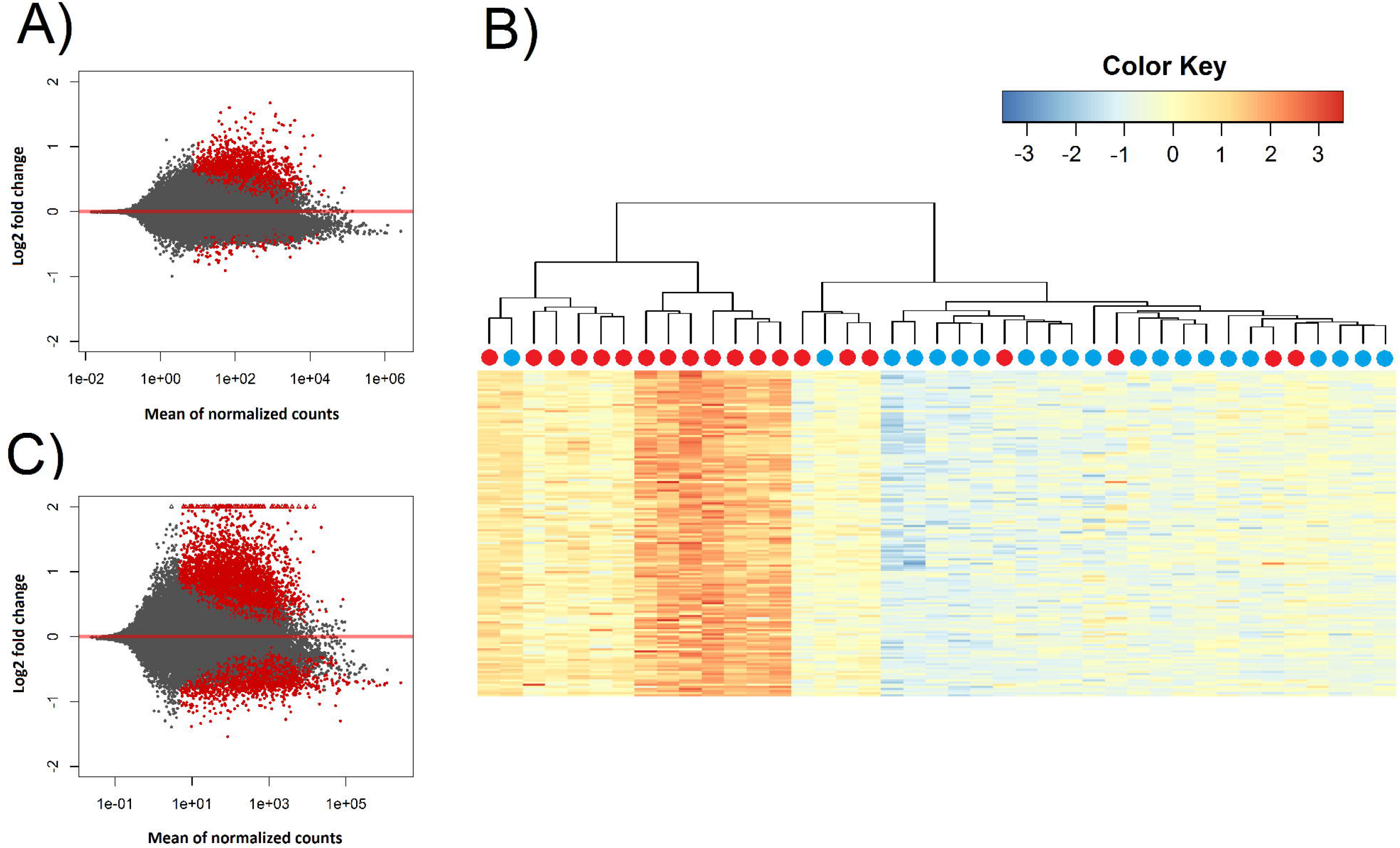
Sample clustering and differential expression. A) Volcano plot showing the result of differential expression analysis between healthy and injured skin (all samples). Level of expression is shown in the x-axis and log_2_ fold change in y-axis, genes with false discovery rate corrected p-values < 0.05 are shown in red. B) Heatmap showing the gene expression of the most significant 156 DE genes (p < 0.001) between healthy (blue) and injured (red) skin. C) Volcano plot showing the result of differential expression analysis between healthy and injured skin using those animals whose injured skin shows some sort of response to *Caligus*. Level of expression is shown in the x-axis and log_2_ fold change in y-axis, genes with false discovery rate corrected p-values < 0.05 are shown in red.

Among these DE genes were well-known components of the innate immune response like interleukins, interferon response factors and complement components (Figure 2A). GO term and KEGG pathway analyses (Additional file 2) revealed a clear enrichment of immune pathways and functions among the up-regulated genes (Figure 2B), highlighting a localised immune response strongly related to cytokines. A similar scenario has been observed in other salmonids such as coho salmon where resistance to sea lice has been associated with early inflammation in skin and head kidney, which results in epithelial hyperplasia and often parasite encapsulation and removal of the sea lice within two weeks [24, 25]. In pink salmon (*Oncorhynchus gorbuscha*), an early and high expression of pro-inflammatory genes (IL-8, TNFα-1, IL-1β) has been suggested as a mechanism of rapid louse rejection [26]. The classical complement pathway has also been linked to resistance of host fish to parasitic copepod infection [7]. The results presented here indicate that despite a marked up-regulation of the local inflammatory response and complement pathway in Atlantic salmon, resembling those of coho salmon or pink salmon, the response does not seem to be sufficient to successfully respond to the louse attachment and feeding.

**Figure 2.**
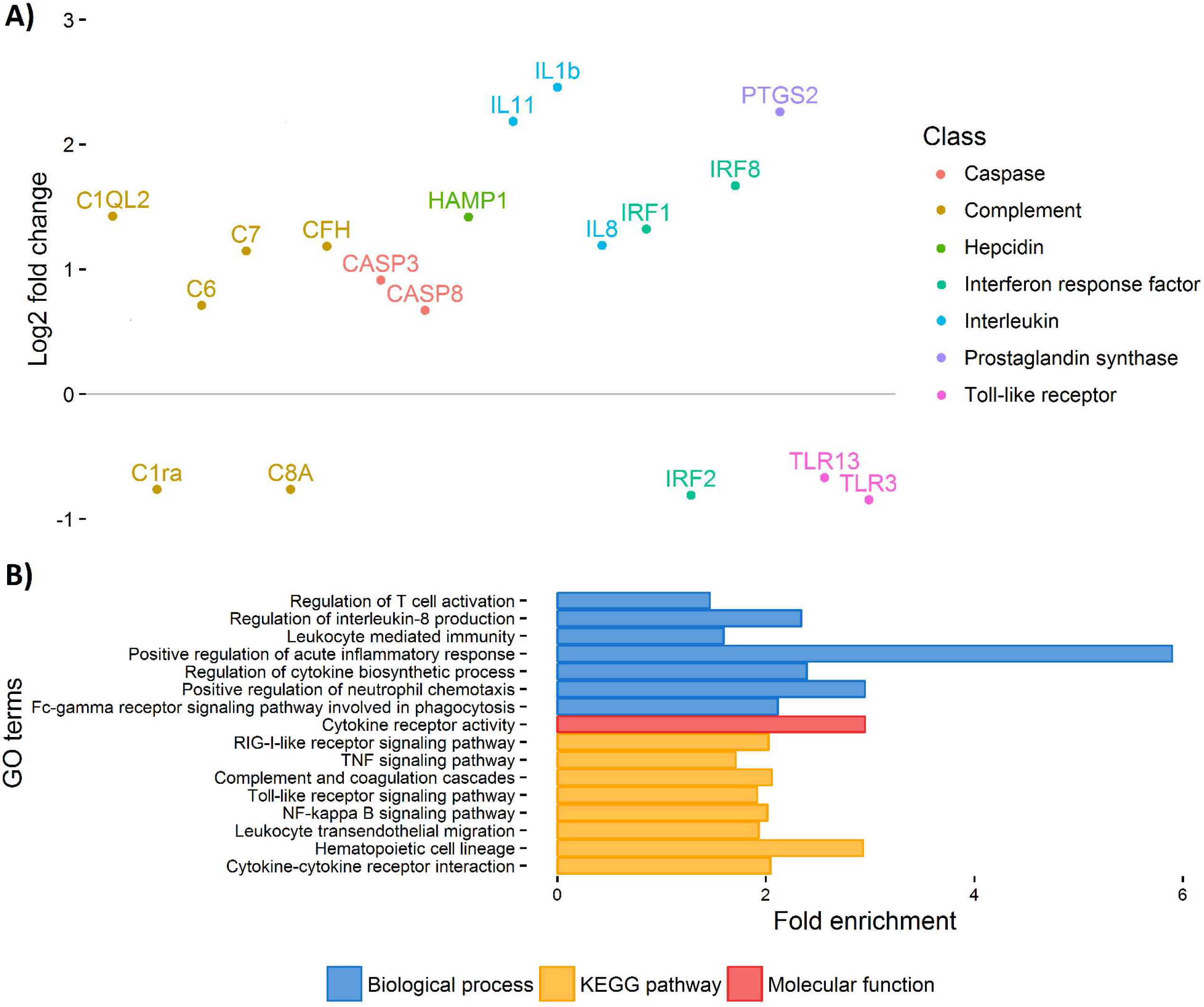
Healthy vs injured skin. A) Important immune-related genes showing differential expression between healthy and injured skin. B) Selection of GO terms enriched amongst DE genes between healthy and injured skin.

In addition to the expected innate immune response observed above, cell division related processes were also clearly up-regulated at louse attachment sites, and well-characterised genes involved in tissue repair such as fibroblast growth factor-binding protein 1 and Epigen showed significant differences between lice attachment sites and healthy skin (FC > 3). Several genes related to the cell matrix and cell adhesion also had higher expression at attachment sites (i.e. cadherin-13, integrin alpha-2, desmoplakin, or various keratin and collagen genes). Cell proliferation is the main response to skin wounds in fish [27], and these results are consistent with those previously found in the early response to *Lepeophtheirus salmonis* [28]. Several mucins were also found to have higher expression at attachment sites, pointing towards increased mucus production and secretion which can also be a typical response to wounding in fish [7].

### Resistance

The Atlantic salmon samples used for RNA-Seq were taken from a broader experiment relating to genetic resistance to sea lice, and were measured for several traits of relevance to resistance and host response to infection. These traits were defined as resistance, measured as number and concentration of lice per fish, and tolerance, defined in this study as weight gain since the start of the challenge. The samples for RNA sequencing were chosen to enable 8 vs 8 comparisons between family-matched fish with high and low values for both resistance (26.2 ± 5.5 vs 54.9 ± 13.5 sea lice per fish) and tolerance (7.0 ± 4.3 vs 28.8 ± 12.3 weight gain percentage).

#### Differential expression

There were 43 genes significantly differentially expressed between resistant and susceptible fish, although all but one were from comparison of healthy skin samples between the two groups (Additional file 3). The susceptible group had higher expression levels for genes involved in muscle contraction like troponins and myosins, which was also highlighted by GO enrichment analyses (Figure 3). Myosins and troponins have previously been identified as genes that respond to sea lice attachment in salmon skin [29]. Further, *Caligus* infection is known to induce increased enzyme activity in muscle tissue [30], and behavioural changes in the fish such as flashing and jumping are associated with ectoparasite removal [31, 32]. It has been recently reported that inactivity or reduced swimming activity contribute to resistance to sea lice [33], so it is possible that the high lice counts of susceptible fish in this study are due to higher activity levels with associated expression of muscle contraction related genes. In turn, high lice burden can provoke behavioural responses increasing fish activity, which results in the up-regulation of muscle genes, increasing the expression differences between resistant-passive-low lice fish and susceptible-active-high lice fish.

**Figure 3.**
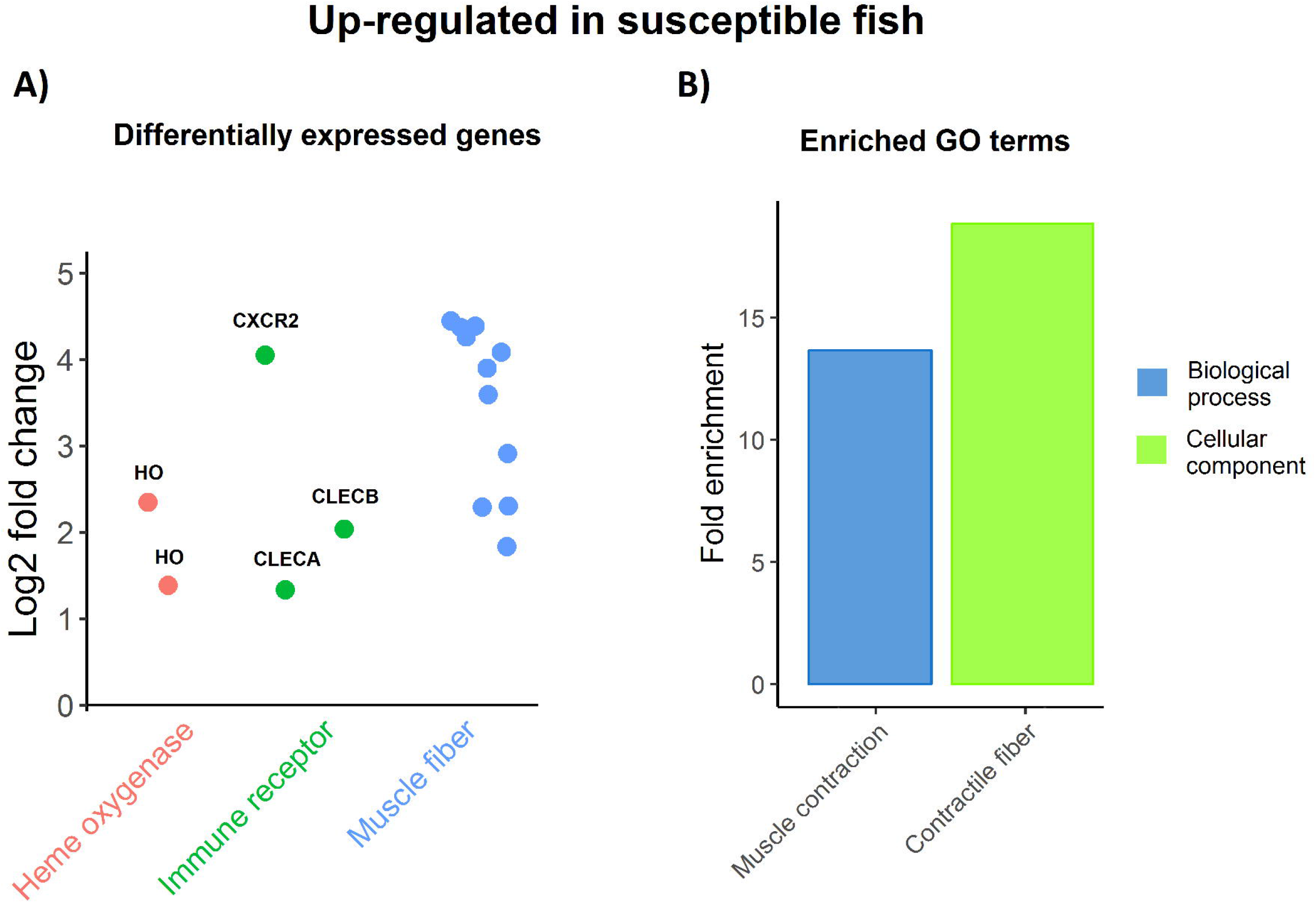
Up-regulation in susceptible fish. A) Genes DE between resistant and susceptible fish, being up-regulated in the latter. B) Enriched GO terms amongst DE genes up-regulated in susceptible fish.

Two heme oxygenase genes (HO), encoding enzymes which catalyze the degradation of heme, also had higher expression levels in susceptible samples (Figure 3). These genes have been previously shown to be up-regulated in response to *Caligus* infection [34]. Reduction of iron availability within the host has been observed in Atlantic salmon infected with *L. Salmonis* [35], as has reduced hematocrit and anemia in Pacific salmon [36]. Finally, three immune receptors showed higher expression in susceptible samples (Figure 3); C-X-C chemokine receptor type 2 (CXCR2) is a receptor for IL-8, its binding causes activation of neutrophils; while C type lectin receptors A and B (CLECA, CLECB) are involved in pathogen recognition and immunity [37]. While it is clear that resistance and host response to sea lice is multifactorial in nature, these genes related to muscle contraction, iron availability and immunity may be targets for functional validation in future studies, and for cross-referencing with genome-wide association analyses to identify candidate causative genes and variants.

#### Network analysis

As an alternative approach to investigate the gene expression profiles associated with resistance and tolerance to sea lice in these samples, a network correlation analysis was performed. For this analysis, instead of categorizing the animals into two groups and testing differential expression, gene expression values were used to group genes with similar expression profiles into different networks. The average expression profile of each network was then correlated with the measured traits to identify significant associations between groups of genes and the resistance phenotypes (Figure 4). One gene network with a total of 76 genes was associated with resistance (as measured by lice counts) in healthy skin (Additional file 3). A single gene from this network was also differentially expressed in the previous analysis between high and low resistant fish, C-type lectin receptor A (FC = -1.34). As previously mentioned, this gene is a receptor involved in antigen recognition and immune response [37]. Four other immune receptors involved in innate immunity and inflammation were assigned to this network and found to be positively correlated with sea lice counts (Table 1). Macrophage mannose receptor 1 (MRC1) shows the highest expression difference between louse attachment vs healthy skin DE genes (FC = 4.79) and also the highest positive correlation with number of sea lice (r = 0.87). MRC1 is also a c-type lectin receptor, expressed in macrophages, dendritic cells and skin in humans. MRC1 plays a role both in innate and adaptive immunity and also acts as a recognition receptor for different pathogens such as bacteria, virus or fungi [38]. Lectins such as MRC1 and CLEC4E have been found to be induced by glucosinolate-enriched feeds in Atlantic salmon, which also reduced lice counts between 17-25% [39]. Lectins have been reported to activate the immune system in response to different parasites in different species [40, 41], therefore modulation of these genes represents a possible route to enhance Atlantic immune responses to sea lice. Two immune receptors were negatively correlated with number of sea lice, CD97 (r = -0.84) and suppressor of cytokine signalling 5 (SOCS5; r = -0.76). CD97 regulates cytokine production and T-cell activation and proliferation [42, 43]; while SOCS5 is part of the cytokine-mediated signalling pathway, and acts as a negative regulator of inflammatory response and other immune-related pathways [44]. The correlation of these genes with number of sea lice is probably indicating that the immune system of the host responds proportionally to the degree of infestation, but nonetheless may be important in the louse-salmon interaction.

**Figure 4.**
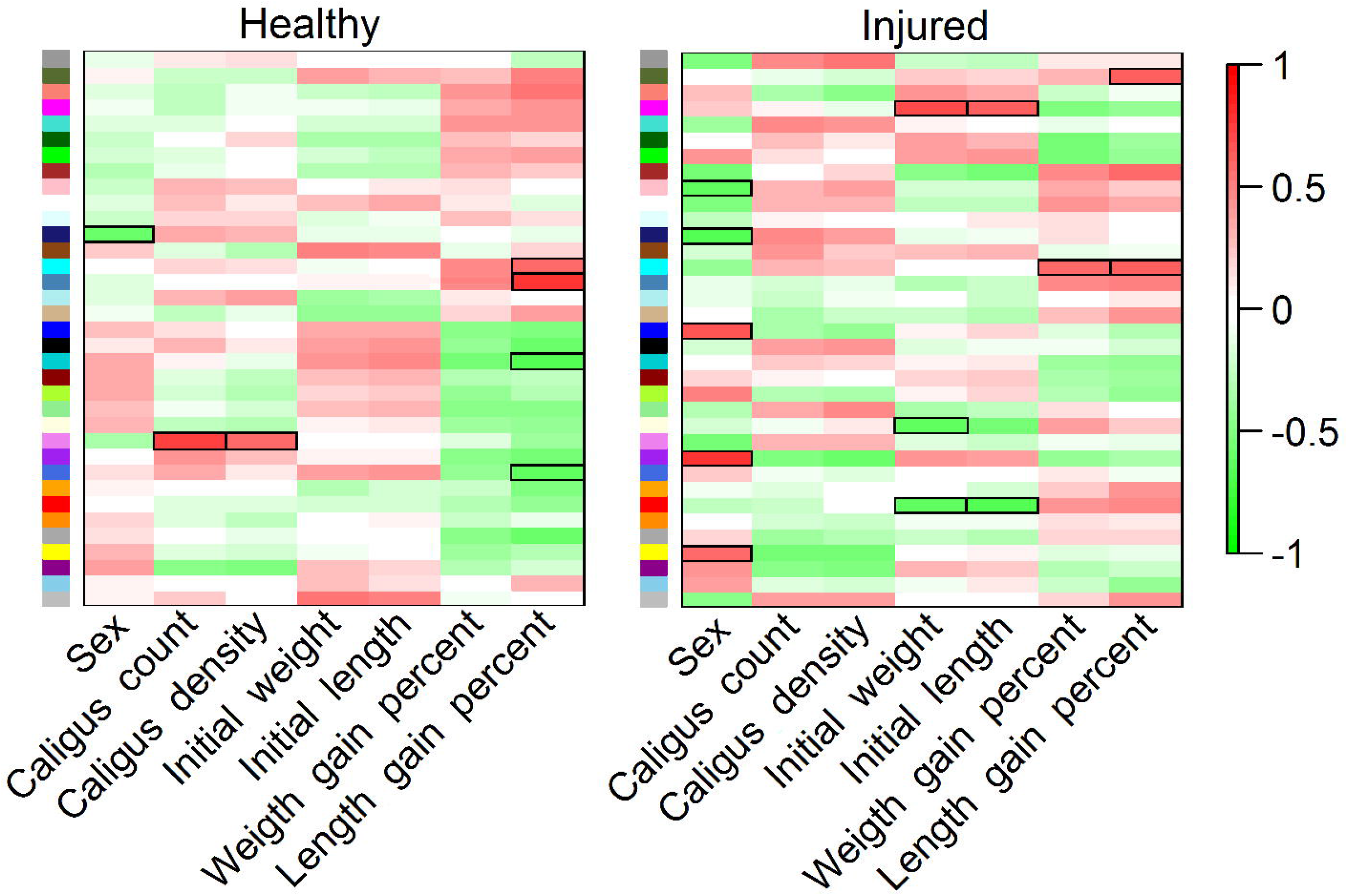
Network correlation analyses. Correlation between gene networks and different traits of interest in healthy and injured skin.

**Table 1.**
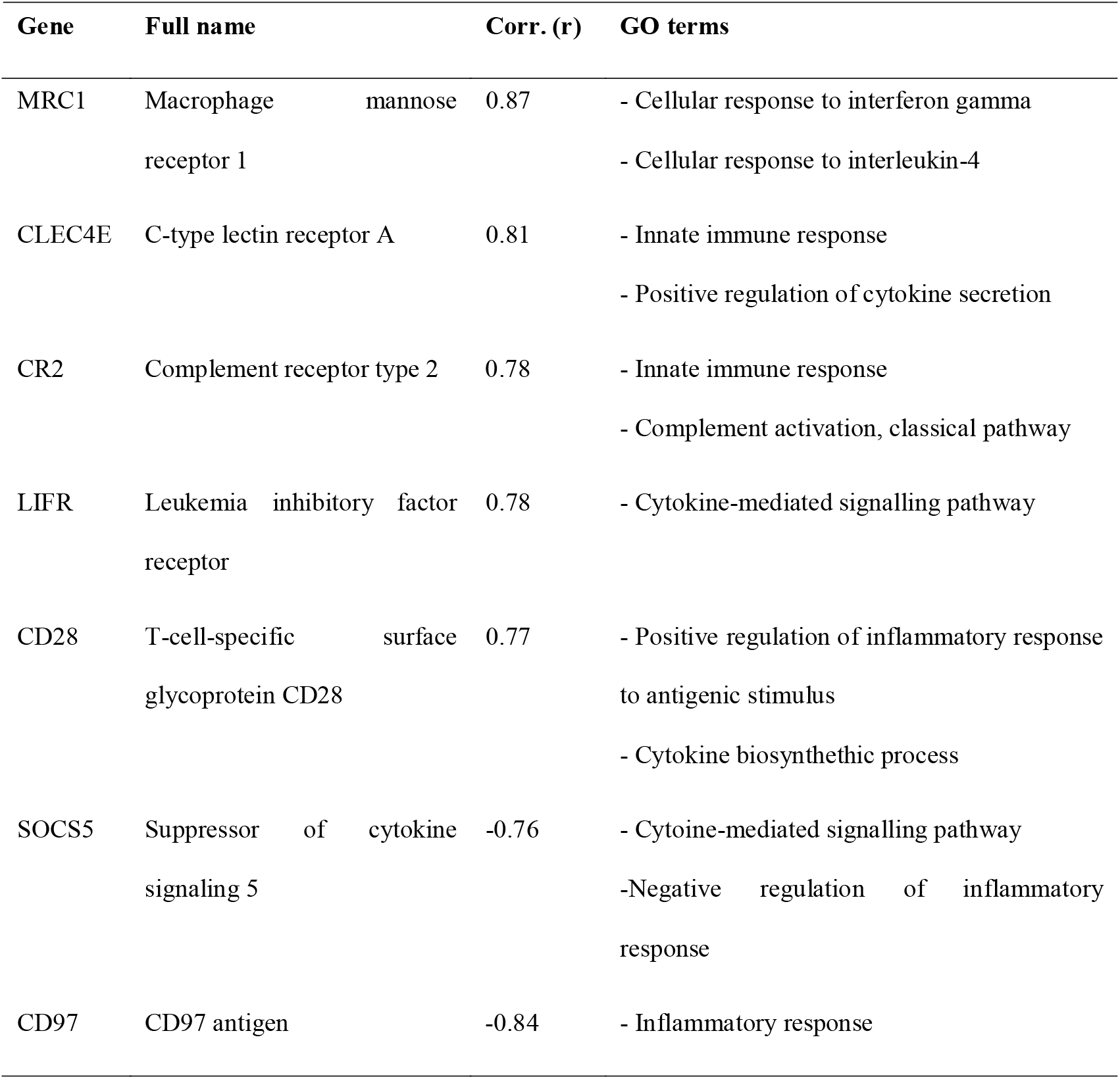
Immune receptors showing correlation with sea lice counts.

Amongst genes without a (well-known) immune function, there was an association between SUMO1 (r = 0.76) and SUMO3 (r = -0.91) expression and resistance. Small Ubiquitin-like Modifier (SUMO) proteins are small proteins similar to ubiquitins that are covalently attached to other proteins to modify their function. According to the gene expression data, SUMO1 seems to be preferred over SUMO3 in salmon upon sea lice infestation. Although post-translational modifications have been barely explored in fish, in mice SUMOylation has been shown to be involved in modulation of host innate immune response to pathogens [45]. SUMOylation is also a very active field of research in plants, where SUMO is known to be involved in many important processes such as plant response to environmental stresses, including pathogens [46]. It would be interesting to further study the role of SUMO in modulating Atlantic salmon responses to sea lice.

### Tolerance

Differences in weight gain percentage from the start to the end of the trial were also investigated; a trait which was defined as tolerance. Tolerance did not show any significant correlation with initial weight (r = -0.27, p = 0.10), sea lice counts (r = 0.12, p = 0.45) or sea lice density (r = 0.19, p = 0.24) in our dataset, and the means for these three traits are not significantly different between our tolerant and not-tolerant groups (t-test p-values > 0.35).

#### Differential expression

A total of 24 and 1 genes were found differentially expressed between fish showing high and low tolerance in healthy and sea louse attachment site samples respectively (Additional file 4). The gene differentially expressed in injured skin, solute carrier family 15 member 1 (SLC15A1), also showed the lowest p-value and highest fold change in healthy skin (FC = 3.38, p = 0.003). SLC15A1 protein is a membrane transporter that mediates the uptake of dipeptides and tripeptides, in humans this gene is expressed in the intestinal epithelium and plays a major role in protein absorption [47]. Another interesting DE gene is myogenic regulatory factor 6 (MYF6; FC = 0.72, p = 0.04). Myogenic regulatory factors are transcription factors that regulate muscle development [48]; in *Senegale sole* decreased expression of these factors was observed in fast muscle when fed with a high-lipid content diet which caused reduced growth [49]. While skin is unlikely to be a highly suitable tissue to study genes underlying fish growth during sea lice infestation, both myogenic factors and increased nutrient absorption, and specifically MYF6 and SLC15A1, are good candidates to better understand growth impairment differences under sea lice infestation.

#### Network analysis

A gene network correlation analysis was also performed for tolerance, finding five associated networks, one of them common between healthy and injured datasets (Figure 4). A total of 373 genes showed correlation values > 0.75 with weight gain percentage (Additional file 4). These genes are fairly heterogeneous, showing no GO or KEGG enrichment. This is perhaps not that surprising; here we have defined tolerance as impact of sea lice on growth performance, but most likely growth performance is only one of the aspects of tolerance. A more tolerant animal will be less affected by the parasite in many ways, and therefore it is logical to find genes involved in different functions correlated with our definition of tolerance. Several immune related genes were positively correlated with weight gain, such as C-X-C chemokine receptor type 4, interferon regulatory factor 2, caspase-8 or TNF receptor-associated factor 5. Very different immune responses for IPNV resistant and IPNV susceptible Atlantic salmon families have been observed [50], so it is well possible that Atlantic salmon can also elicit different immune responses against sea lice, leading to different outcomes. However, while IPNV resistance is mainly explained by a single locus [51], sea lice tolerance is most likely highly polygenic, with probably different mechanisms operating in the different families, which hinders the understanding of tolerance (and resistance) mechanisms.

### Signatures of differentiation

A total of 126,741 SNPs were identified from the RNA-Seq of the 21 individual fish. These were used to compute Fst values between the resistant and susceptible samples, and between tolerant and not tolerant samples, through a sliding window approach (100 Kb windows every 50 Kb). Only windows with ≥ 5 SNPs were retained for further analysis. Genomic windows with Fst values ≥ 0.20 were considered to show putative signs of differentiation between both groups (35 out of 15,321 total windows, 99.8 percentile). 35 differential genomic regions were found to be associated with resistance and 28 with tolerance (Additional file 5). While the tolerance results did not overlap with differentially expressed genes, genomic regions showing signs of differentiation between resistant and susceptible sample did highlight some interesting candidate genes (Figure 5). The genomic region showing the clearest sign of differentiation between resistant and susceptible samples was found in chromosome 5 between 18.50 and 18.75 Mb, with four consecutive windows with Fst values between 0.237 and 0.306. There are 29 SNPs and 7 genes in this genomic region (Figure 5B). One of these genes, Neurexophilin And PC-Esterase Domain Family Member 3 (NXPE3), was also differentially expressed between resistant and susceptible samples (p-val = 0.02, FC = -0.92). One of the SNPs in the region (Fst = 0.333) codes for a non-synonymous mutation in NXPE3, and four additional SNPs (Fst = 0.421-0.454) could fall within coding regions of NXPE3 as well (these SNPs fall in a transcribed region without annotation in the official salmon annotation; blast searches against NCBI’s nr database match NXPE3). Despite being conserved in vertebrates there is not much information about this gene. In humans it binds alpha neurexins, a group of presynaptic transmembrane receptors that promote adhesion between dendrites and axons [52]. Neurexins have been linked to behavioural response to parasites in bees [53] and in host response to parasite infection in hard clam, *Mercenaria mercenaria* [54]. While not an obvious functional candidate, NXPE3 is worthy of further investigation in host resistance to sea lice due to its differential expression and overlap with a genomic region showing putative differentiation between resistant and susceptible fish. Another interesting gene in the same region of chromosome 5 is COP9 signalosome complex subunit 6 (COPS6). This gene is highly expressed in our samples and contains three SNPs, the one with the highest Fst (0.312) codes for a synonymous mutation. This gene is part of the COP9 signalosome complex, which is a regulator of the ubiquitin conjugation pathway but is also involved in phosphorylation-mediated regulation of various immune-related genes, such as NF-kappa-B-inhibitor alpha or IRF8.

**Figure 5.**
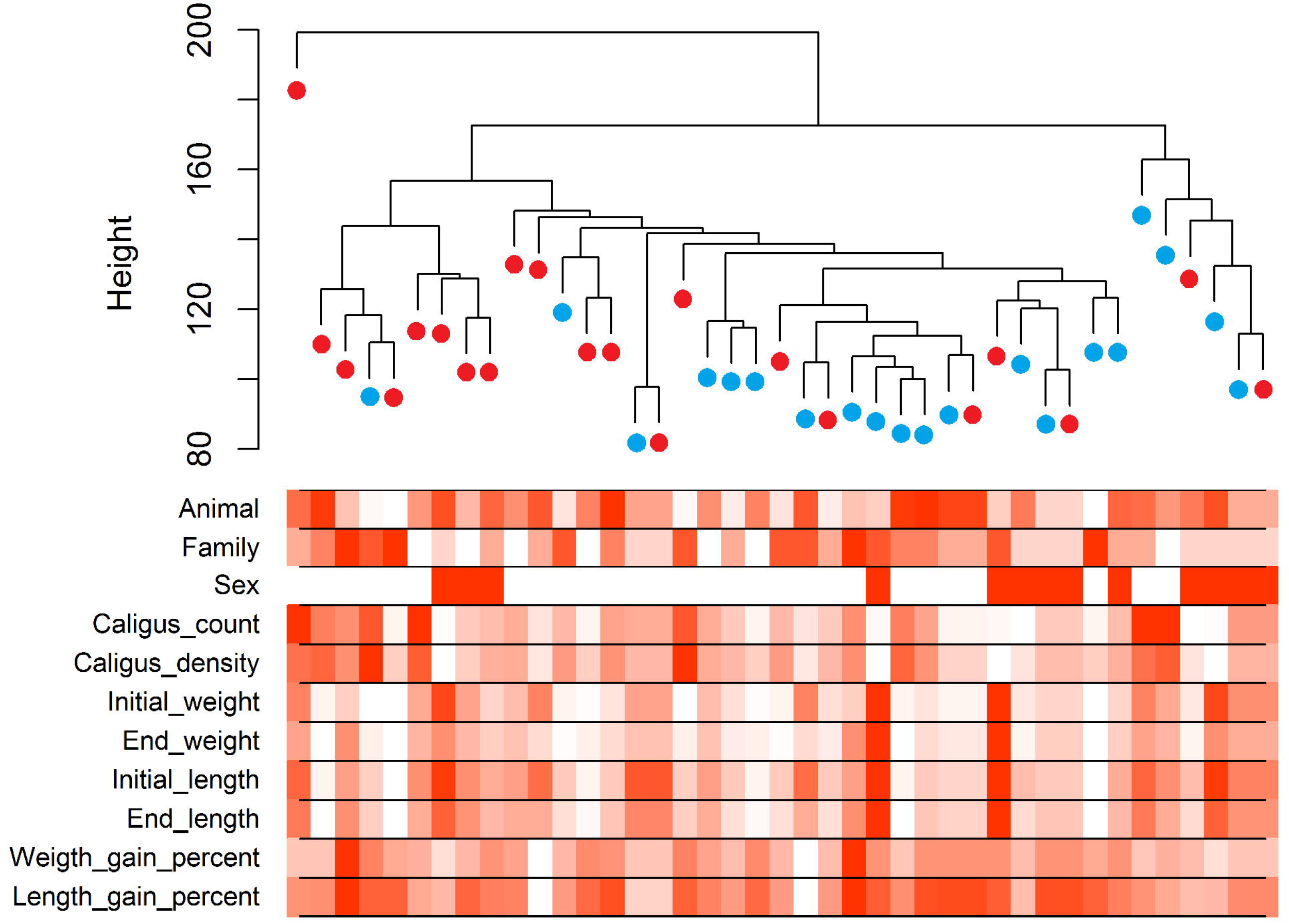
Fst between resistant and susceptible fish. A) Manhattan plot showing the chromosomes and position in the x-axis and Fst in the y-axis. B) High Fst region in chromosome 5.

## CONCLUSIONS

The results of this study highlight that the early gene expression response of Atlantic salmon to sea lice involves up-regulation of many different components of the immune system (inflammatory response, cytokine production, TNF and NF-kappa B signalling and complement activation) along with tissue repair activation. The comparison of resistant versus susceptible animals highlighted enrichment of pathways related to fish activity, iron availability and receptors modulating pathogen recognition and immune response, and identified a few candidate genomic regions potentially involved in genetic resistance. Overall, this study contributes to an improved understanding of Atlantic salmon early response to sea lice in skin, and into the gene expression profiles underpinning genetic resistance to sea lice in salmon. The identified pathways and genes may be targets for future studies aimed at development of new treatments, vaccines or prevention strategies. The data can also be cross-referenced with high power genome-wide association studies to help prioritise putative causative genes and variants that have potential to improve genomic selection programs for genetic improvement of resistance to this industry’s most serious disease.

## METHODS

### Experimental design

2,668 Atlantic salmon (*Salmo salar*) pre-smolts (average weight 136 g) from 104 families from the breeding population of Salmones Chaicas, Xth Region, Chile, were experimentally challenged with *Caligus rogercresseyi* (chalimus II-III). Briefly, infestation with the parasite was carried out by using 13 to 24 copepodids per fish and stopping the water flow for 6 h after infestation. Eight days after the infestation fish were euthanized and fins from each fish were collected and fixed for processing and lice counting.

42 samples from 21 fish from six different families (2 to 5 fish per family) were selected for RNA sequencing (Additional file 6) based on the traits of interest (number of sea lice attached to their fins and growth during challenge). Skin samples (both from attachment sites and health skin) were obtained from each animal and stored in RNAlater at 4 °C for 24 h, and then at -20°C until RNA extraction for sequencing.

### RNA extraction and sequencing

For all the 42 samples a standard TRI Reagent RNA extraction protocol was followed. Briefly, approximately 50 mg of skin was homogenized in 1 ml of TRI Reagent (Sigma, St. Louis, MO) by shaking using 1.4 mm silica beads, then 100 μl of 1-bromo-3-chloropropane (BCP) was added for phase separation. This was followed by precipitation with 500 μl of isopropanol and posterior washes with 65-75 % ethanol. The RNA was then resuspended in RNAse-free water and treated with Turbo DNAse (Ambion). Samples were then cleaned up using Qiagen RNeasy Mini kit columns and their integrity was checked on Agilent 2200 Bioanalyzer (Agilent Technologies, USA). Thereafter, the Illumina Truseq mRNA stranded RNA-Seq Library Prep Kit protocol was followed directly. Libraries were checked for quality and quantified using the Bioanalyzer 2100 (Agilent), before being sequenced on three lanes of the Illumina Hiseq 4000 instrument using 75 base paired-end sequencing at Edinburgh Genomics, UK. Raw reads have been deposited in NCBI’s Sequence Read Archive (SRA) under accession number SRP100978.

### Read mapping

The quality of the sequencing output was assessed using FastQC v.0.11.5 (http://www.bioinformatics.babraham.ac.uk/projects/fastqc/). Quality filtering and removal of residual adaptor sequences was conducted on read pairs using Trimmomatic v.0.32 [55]. Specifically, Illumina specific adaptors were clipped from the reads, leading and trailing bases with a Phred score less than 20 were removed and the read trimmed if the sliding window average Phred score over four bases was less than 20. Only reads where both pairs were longer than 36 bp post-filtering were retained. Filtered reads were mapped to the most recent Atlantic salmon genome assembly (ICSASG_v2; Genbank accession GCF_000233375.1 [23]) using STAR v.2.5.2b [56], the maximum number of mistmatches for each read pair was set to 10 % of trimmed read length, and minimum and maximum intron lengths were set to 20 bases and 1 Mb respectively. Uniquely mapped paired-reads were counted and assigned to genes (NCBI *Salmo salar* Annotation Release 100) using FeatureCounts [57], included in the SourceForge Subread package v.1.5.0. Only reads with both ends mapped to the same gene were considered in downstream analyses.

### SNP identification

SNPs were identified and genotypes for individual samples were called using samtools v1.2 [58]. Reads with mapping quality < 30 and bases with phred quality scores < 30 were excluded. To call a SNP, a read depth ≥ 10 and ≥ 3 reads for the alternative allele were required. Only SNPs called in > 75% of the samples were retained for downstream analyses. The putative effect of the SNPs was assessed using the official salmon genome annotation (NCBI *Salmo salar* Annotation Release 100) and the SnpEff v.4.2 software [59].

### Differential Expression

Statistical analyses related to differential expression and pathway enrichment were performed using R v.3.3.1 [60]. Gene count data were used to estimate differential gene expression using the Bioconductor package DESeq2 v.3.4 [61]. The Benjamini-Hochberg false discovery rate (FDR) was applied, and transcripts with corrected p-values < 0.05 and absolute log_2_ fold change values (FC) > 0.5 were considered differentially expressed genes. Samples were hierarchically clustered according to gene read counts after a variance stabilizing transformation, using Euclidean as the distance measure and complete-linkage as the agglomeration method (R package flashClust [62]). Heatmaps of gene expression were created using the R package gplots v3.0.1 heatmap.2 function, using read counts after regularized log transformation (DESeq2 [61]).

### Pathway Enrichment

Gene Ontology (GO) enrichment analyses were performed using Blast2GO v.4.1 [63]. Briefly, genes showing > 10 reads in > 90 % of the samples were annotated against the manually curated protein database Swiss-Prot [64] and GO terms were assigned to them using Blast2GO. GO enrichment for specific genes lists was tested against the whole set of expressed genes using Fisher’s Exact Test. GO terms with ≥ 5 DE genes assigned and showing a Benjamini-Hochberg FDR corrected p-value < 0.05 were considered enriched. Kyoto Encyclopedia of Genes and Genomes (KEGG) enrichment analyses were performed using KOBAS v3.0.3 [65]. Briefly, genes showing > 10 reads in > 90 % of the samples were annotated against KEGG protein database [66] to determine KEGG Orthology (KO). KEGG enrichment for specific gene lists was tested by comparison to the whole set of expressed genes using Fisher’s Exact Test. KEGG pathways with ≥ 5 DE genes assigned and showing a Benjamini-Hochberg FDR corrected p-value < 0.05 were considered enriched.

### Network Correlation Analysis

Gene expression network correlation analyses were performed using the R package WGCNA [67]. Read counts after variance stabilizing transformation were used as a measure of gene expression. The genes were clustered using a power of 6 (scale free topology index > 0.9), a minimum of 30 genes per cluster and merging those clusters with a distance < 0.25 between them. Correlation between network summary profiles and external traits was quantified, and network-trait association showing p-values < 0.05 were considered significant. Genes showing network membership significance values < 0.05 and | r | > 0.75 were considered to be correlated with the trait of interest.

### Fst Analyses

Weir and Cockerham’s Fst statistics were calculated using VCFtools v0.1.14 [68]. Mean Fst values were calculated using a sliding window approach with 100 Kb windows and a step of 50 Kb. Only those windows containing at least 5 SNPs were considered.

## DECLARATIONS

### Acknowledgements

The authors would like to thank the contribution of Aquainnovo and Salmones Chaicas for providing the biological material and phenotypic records of the experimental challenges.

### Authors contributions

RDH, JMY and DR were responsible for the concept and design of this work. AB managed the collection of the samples. AP performed the molecular biology experiments. DR performed bioinformatic and statistical analyses. DR, RDH and JMY drafted the manuscript. All authors read and approved the final manuscript.

### Ethics approval and consent to participate

The lice challenge experiments were performed under local and national regulatory systems and were approved by the Animal Bioethics Committee (ABC) of the Faculty of Veterinary and Animal Sciences of the University of Chile (Santiago, Chile), Certificate N° 01-2016, which based its decision on the Council for International Organizations of Medical Sciences (CIOMS) standards, in accordance with the Chilean standard NCh-324-2011.

### Consent to publish

Not applicable

### Availability of data and materials

The datasets generated during the current study are available in the NCBI’s Sequence Read Archive (SRA) repository under accession number SRP100978 [https://www.ncbi.nlm.nih.gov/sra/?term=SRP100978].

## ADDITIONAL FILES

### Additional file 1

PNG (.png)

Hierarchical cluster of the whole dataset.

Hierarchical clustering for all the samples using the expression of all the genes in our dataset. Healthy (blue) and injured (red) sample phenotypes are shown using a white (minimum) to red (maximum) color code.

### Additional file 2

Excel (.xlsx)

Comparison between healthy and injured skin.

Differentially expressed genes (1^st^ sheet), GO term and KEGG pathway enrichment (2^nd^ sheet) are shown for the comparison between healthy and injured skin.

### Additional file 3

Excel (.xlsx)

Comparison between resistant and susceptible samples.

Differentially expressed genes (1^st^ sheet), GO term and KEGG pathway enrichment (2^nd^ sheet), and network analysis results (3^rd^ sheet) are shown for the comparison between resistance and susceptible samples.

### Additional file 4

Excel (.xlsx)

Comparison between tolerant and not tolerant samples.

Differentially expressed genes (1^st^ sheet) and network analysis (2^nd^ sheet) are shown for the comparison between tolerant and not tolerant samples.

### Additional file 5

Excel (.xlsx)

Fst values for resistance and tolerance comparisons.

Genomic regions showing Fst values > 1 for resistance (1^st^ sheet) and tolerance (2^nd^ sheet) comparisons.

### Additional file 6

Excel (.xlsx)

RNA-seq samples

List of samples sequenced in this study and their phenotypes.

